## Supplementary material for "Unified inference of missense variant effects and gene constraints in the human genome"

Yi-Fei Huang

Department of Biology and Huck Institutes of the Life Sciences,
Pennsylvania State University, University Park, PA 16802, USA

### Supplemental Table S1: Genomic features for UNEECON.

| Feature group | Feature name | Type | Reference | Note |
|---|---|---|---|---|
| Sequence conservation | SIFT prediction | Binary | [1] | Binary prediction of deleteriousness from SIFT |
| | LRT prediction | Binary | [1] | Binary prediction of deleteriousness from LRT |
| | MA prediction | Binary | [1] | Binary prediction of deleteriousness from Mutation Assessor |
| | PROVEAN prediction | Binary | [1] | Binary prediction of deleteriousness from PROVEAN |
| | SLR score | Binary | [1] | Raw SLR score |
| | SIFT score | Numeric | [1] | Raw SIFT score |
| | LRT omega | Numeric | [1] | Raw LRT score |
| | MA score | Numeric | [1] | Raw Mutation Assessor score |
| | PROVEAN score | Numeric | [1] | Raw PROVEAN score |
| | Grantham score | Numeric | [2] | Raw Grantham score |
| | HMM entropy | Numeric | [3] | HMM entropy score from SNVBox |
| | HMM relative entropy | Numeric | [3] | HMM relative entropy score from SNVBox |
| | dscore | Numeric | [1] | Dscore from PolyPhen-2 |
| | Primate phyloP score | Numeric | [4] | Primate phyloP conservation score |
| | Mammalian phyloP score | Numeric | [4] | Mammalian phyloP conservation score |
| | Vertebrate phyloP score | Numeric | [4] | Vertebrate phyloP conservation score |
| Structural information | PredRSAB | Numeric | [3] | Probability of the residue being buried |
| | PredRSAI | Numeric | [3] | Probability of the residue being intermediately exposed |
| | PredRSAE | Numeric | [3] | Probability of the residue being exposed |
| | PredBFactorF | Numeric | [3] | Probability that the residue's backbone is flexible |
| | PredBFactorM | Numeric | [3] | Probability that the residue's backbone is intermediately flexible |
| | PredBFactorS | Numeric | [3] | Probability that the residue's backbone is stiff |
| | PredStabilityH | Numeric | [3] | Probability that the residue strongly stabilizes folding |
| | PredStabilityM | Numeric | [3] | Probability that the residue stabilizes folding |
| | PredStabilityL | Numeric | [3] | Probability that the residue destabilizes folding |
| | PredSSE | Numeric | [3] | Probability that the secondary structure of the residue is strand |
| | PredSSH | Numeric | [3] | Probability that the secondary structure of the residue is helix |
| | PredSSC | Numeric | [3] | Probability that the secondary structure of the residue is loop |
| Regulatory information | SPIDEX | Numeric | [5] | SPIDEX Splicing score |
| | Maximum RNA-seq signal | Numeric | [6] | Maximum RNA-seq signal from the Roadmap Epigenomics Project |

Supplemental Table S2: Statistical significance of the difference of AUCs between UNEECON and alternative methods in predicting ClinVar missense variants associated with autosomal dominant disorders.

| UNEECON | Predicting pathogenic missense variants labeled as "autosomal dominant" in ClinVar | Predicting pathogenic missense variants within autosomal dominant genes |
|---|---|---|
| *vs* MPC | 0.04618 * | 3.190e-12 ** |
| *vs* LASSIE | 0.0001567 ** | 0.2367 |
| *vs* PrimateAI | 0.04633 * | 8.233e-11 ** |
| *vs* Eigen | 0.001115 ** | 3.004e-05 ** |
| *vs* CADD | 4.869e-05 ** | 1.077e-08 ** |
| *vs* RVIS | 3.155e-61 ** | 1.468e-96 ** |
| *vs* pLI | 5.414e-51 ** | 5.065e-108 ** |
| *vs* CCR | 2.082e-19 ** | 1.832e-60 ** |

The numbers represent $p$-values from the DeLong test [7]. **: $p$-value $< 0.01$; *: $p$-value $< 0.05$.

Supplemental Table S3: Statistical significance of the difference of AUCs between UNEECON-G and alternative methods in predicting disease genes and essential genes.

| UNEECON-G | HI gene | Autosomal dominant gene | MGI essential gene | CRISPR essential gene |
|---|---|---|---|---|
| *vs* pLI | 0.4504 | 5.541e-09** | 5.153e-08** | 3.893e-19** |
| *vs* mis-z | 3.514e-8** | 2.530e-12** | 1.449e-28** | 0.0001982** |
| *vs* RVIS | 1.007e-12** | 1.852e-12** | 1.969e-25** | 0.0002210** |
| *vs* GDI | 1.930e-08** | 4.220e-12** | 3.719e-39** | 1.043e-26** |

The numbers represent $p$-values from the DeLong test [7]. **: $p$-value $< 0.01$; *: $p$-value $< 0.05$.

Supplemental Table S4: Enrichment of Reactome pathways in the 956 genes intolerant to both missense and loss-of-function mutations. The 956 genes tolerant to missense but not to loss-of-function mutations are utilized as the background gene set.

| Category | Fold enrichment | *p*-value | FDR |
|---|---|---|---|
| Opioid Signalling (R-HSA-111885) | 16.95 | 1.36E-04 | 1.60E-02 |
| Neurotransmitter receptors and postsynaptic signal transmission (R-HSA-112314) | 11.96 | 2.72E-08 | 1.04E-05 |
| Transmission across Chemical Synapses (R-HSA-112315) | 8.47 | 3.32E-10 | 2.54E-07 |
| Neuronal System (R-HSA-112316) | 4.20 | 1.58E-10 | 2.42E-07 |
| mRNA Splicing (R-HSA-72172) | 3.99 | 2.25E-06 | 5.75E-04 |
| mRNA Splicing - Major Pathway (R-HSA-72163) | 3.99 | 2.25E-06 | 4.92E-04 |
| G2/M Transition (R-HSA-69275) | 3.29 | 5.27E-04 | 4.75E-02 |
| Processing of Capped Intron-Containing Pre-mRNA (R-HSA-72203) | 2.83 | 3.63E-05 | 5.05E-03 |
| Innate Immune System (R-HSA-168249) | 2.21 | 2.56E-05 | 3.92E-03 |
| Metabolism of RNA (R-HSA-8953854) | 2.12 | 4.32E-06 | 8.27E-04 |
| Developmental Biology (R-HSA-1266738) | 1.85 | 7.35E-06 | 1.25E-03 |
| Axon guidance (R-HSA-422475) | 1.85 | 3.58E-04 | 3.65E-02 |
| Metabolism of proteins (R-HSA-392499) | 1.77 | 3.51E-07 | 1.08E-04 |
| Post-translational protein modification (R-HSA-597592) | 1.68 | 1.48E-04 | 1.61E-02 |
| Metabolism (R-HSA-1430728) | 1.64 | 5.19E-04 | 4.97E-02 |
| Signal Transduction (R-HSA-162582) | 1.46 | 3.63E-05 | 4.63E-03 |
| Unclassified (UNCLASSIFIED) | .68 | 1.63E-09 | 8.34E-07 |

Supplemental Table S5: Enrichment of Gene Ontology (molecular function) terms in the 956 genes intolerant to both missense and loss-of-function mutations. The 956 genes tolerant to missense but not to loss-of-function mutations are utilized as the background gene set.

| Category | Fold enrichment | *p*-value | FDR |
|---|---|---|---|
| potassium channel activity (GO:0005267) | 16.95 | 1.36E-04 | 4.29E-03 |
| potassium ion transmembrane transporter activity (GO:0015079) | 5.23 | 8.46E-04 | 2.04E-02 |
| ligand-gated channel activity (GO:0022834) | 4.19 | 2.34E-03 | 4.35E-02 |
| ligand-gated ion channel activity (GO:0015276) | 4.19 | 2.34E-03 | 4.16E-02 |
| GTPase activity (GO:0003924) | 3.49 | 2.09E-04 | 6.13E-03 |
| ion transmembrane transporter activity (GO:0015075) | 3.19 | 2.81E-05 | 1.44E-03 |
| cation transmembrane transporter activity (GO:0008324) | 3.06 | 1.23E-04 | 4.58E-03 |
| inorganic cation transmembrane transporter activity (GO:0022890) | 3.06 | 1.23E-04 | 4.20E-03 |
| pyrophosphatase activity (GO:0016462) | 2.76 | 1.44E-05 | 1.47E-03 |
| nucleoside-triphosphatase activity (GO:0017111) | 2.76 | 1.44E-05 | 1.18E-03 |
| hydrolase activity, acting on acid anhydrides, in phosphorus-containing anhydrides (GO:0016818) | 2.76 | 1.44E-05 | 9.83E-04 |
| hydrolase activity, acting on acid anhydrides (GO:0016817) | 2.76 | 1.44E-05 | 8.43E-04 |
| mRNA binding (GO:0003729) | 2.68 | 1.83E-03 | 3.76E-02 |
| hydrolase activity (GO:0016787) | 2.17 | 1.53E-07 | 2.09E-05 |
| transmembrane transporter activity (GO:0022857) | 2.13 | 4.26E-04 | 1.16E-02 |
| transporter activity (GO:0005215) | 2.12 | 5.97E-05 | 2.45E-03 |
| RNA binding (GO:0003723) | 1.94 | 5.96E-04 | 1.53E-02 |
| protein kinase activity (GO:0004672) | 1.91 | 9.95E-04 | 2.27E-02 |
| phosphotransferase activity, alcohol group as acceptor (GO:0016773) | 1.87 | 9.96E-04 | 2.15E-02 |
| catalytic activity (GO:0003824) | 1.81 | 1.27E-12 | 5.20E-10 |
| transferase activity, transferring phosphorus-containing groups (GO:0016772) | 1.73 | 2.24E-03 | 4.37E-02 |
| transferase activity (GO:0016740) | 1.73 | 2.83E-05 | 1.29E-03 |
| Unclassified (UNCLASSIFIED) | .69 | 6.78E-11 | 1.39E-08 |

Supplemental Table S6: Enrichment of Gene Ontology (biological process) terms in the 956 genes intolerant to both missense and loss-of-function mutations. The 956 genes tolerant to missense but not to loss-of-function mutations are utilized as the background gene set.

| Category | Fold enrichment | *p*-value | FDR |
|---|---|---|---|
| regulation of membrane potential (GO:0042391) | 6.73 | 2.96E-05 | 5.25E-03 |
| RNA splicing (GO:0008380) | 5.13 | 7.19E-06 | 8.91E-03 |
| mRNA splicing, via spliceosome (GO:0000398) | 4.98 | 1.23E-05 | 7.62E-03 |
| RNA splicing, via transesterification reactions with bulged adenosine as nucleophile (GO:0000377) | 4.98 | 1.23E-05 | 5.08E-03 |
| RNA splicing, via transesterification reactions (GO:0000375) | 4.98 | 1.23E-05 | 3.81E-03 |
| RNA processing (GO:0006396) | 3.57 | 2.56E-05 | 6.34E-03 |
| trans-synaptic signaling (GO:0099537) | 3.41 | 6.53E-05 | 9.00E-03 |
| synaptic signaling (GO:0099536) | 3.41 | 6.53E-05 | 8.10E-03 |
| anterograde trans-synaptic signaling (GO:0098916) | 3.32 | 1.03E-04 | 1.07E-02 |
| chemical synaptic transmission (GO:0007268) | 3.32 | 1.03E-04 | 9.87E-03 |
| regulation of biological quality (GO:0065008) | 2.28 | 4.91E-04 | 4.35E-02 |
| intracellular signal transduction (GO:0035556) | 2.20 | 6.70E-05 | 7.55E-03 |
| signal transduction (GO:0007165) | 1.75 | 3.00E-05 | 4.64E-03 |
| cellular response to stimulus (GO:0051716) | 1.66 | 2.64E-05 | 5.45E-03 |

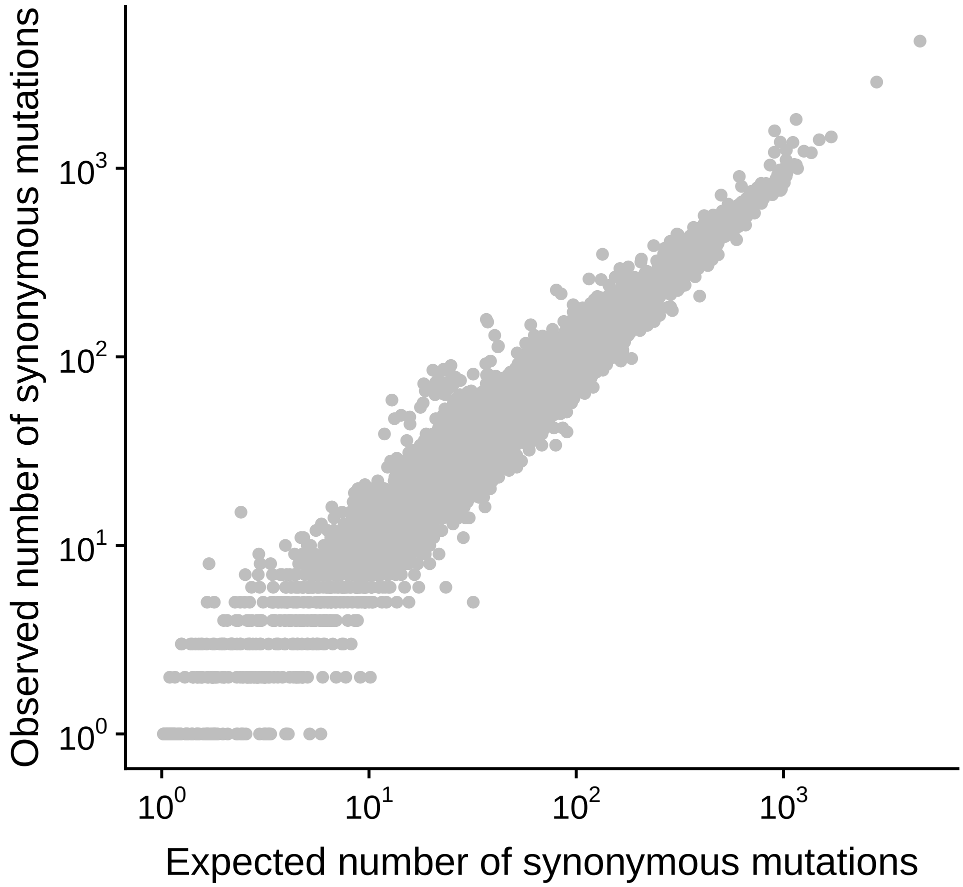


Supplemental Figure S1: Correlation between the expected and the obeserved numbers of synonymous mutations across protein-coding genes under the neutral mutation model. The observed number of synonymous mutations for each gene is derived from the gnomAD exome sequencing data. The expected number of synonymous mutations is predicted by UNEECON's context-dependent mutation model.

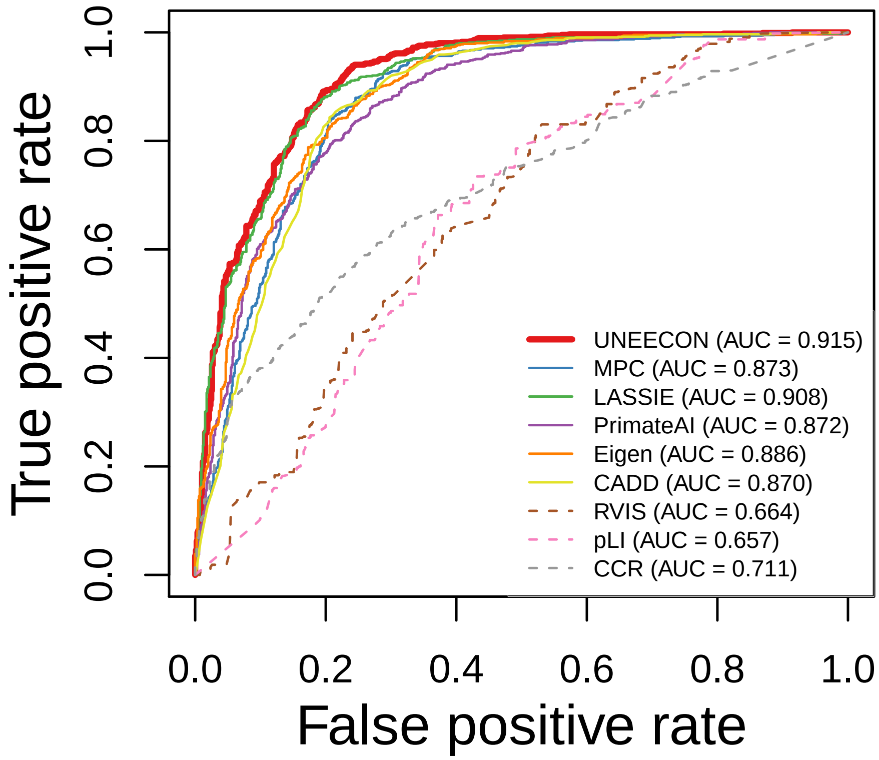


Supplemental Figure S2: Performance of UNEECON and alternative methods in predicting ClinVar pathogenic variants within autosomal dominant genes. Benign missense variants from ClinVar are utilized as negative controls.

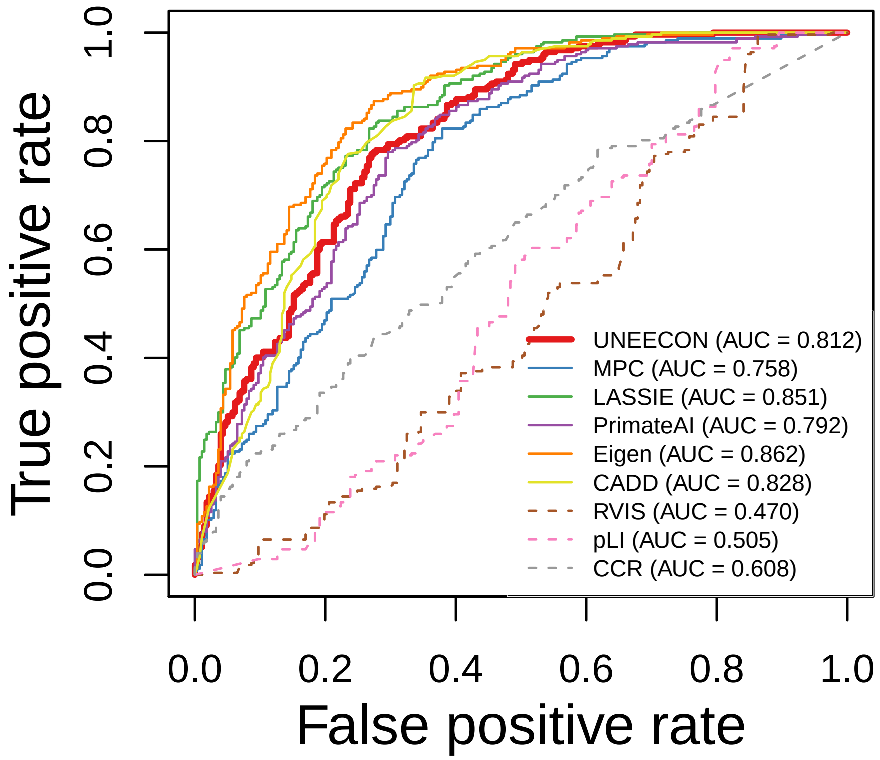


Supplemental Figure S3: Performance of UNEECON and alternative methods in predicting CinVar pathogenic variants with an autosomal recessive mode of inheritance. Benign missense variants from ClinVar are utilized as negative controls.

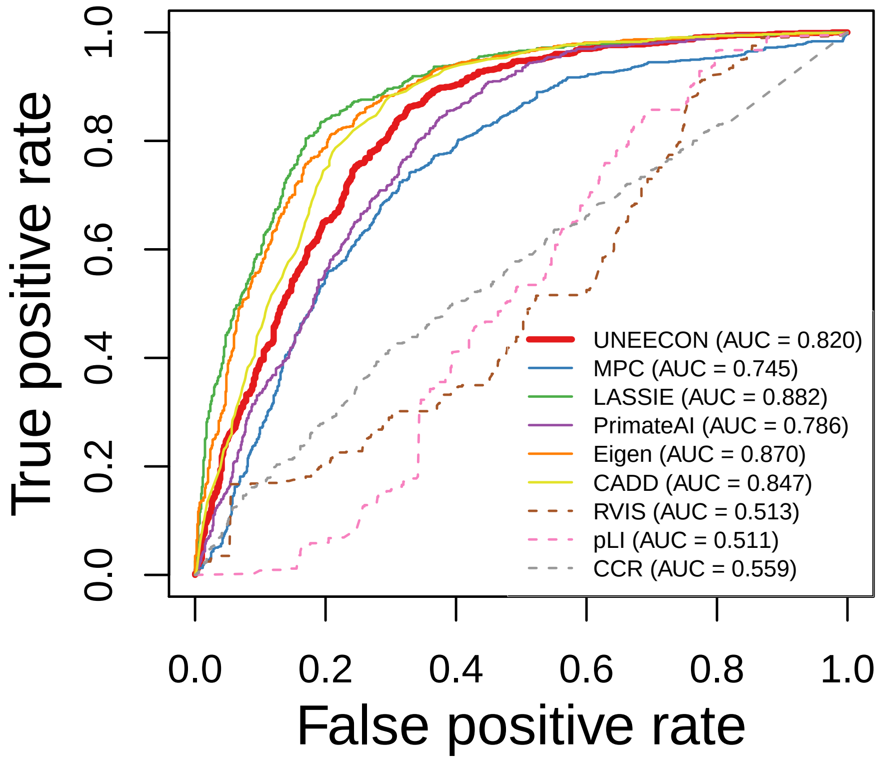


Supplemental Figure S4: Performance of UNEECON and alternative methods in predicting ClinVar pathogenic variants within autosomal recessive disease genes. Benign missense variants from ClinVar are utilized as negative controls.

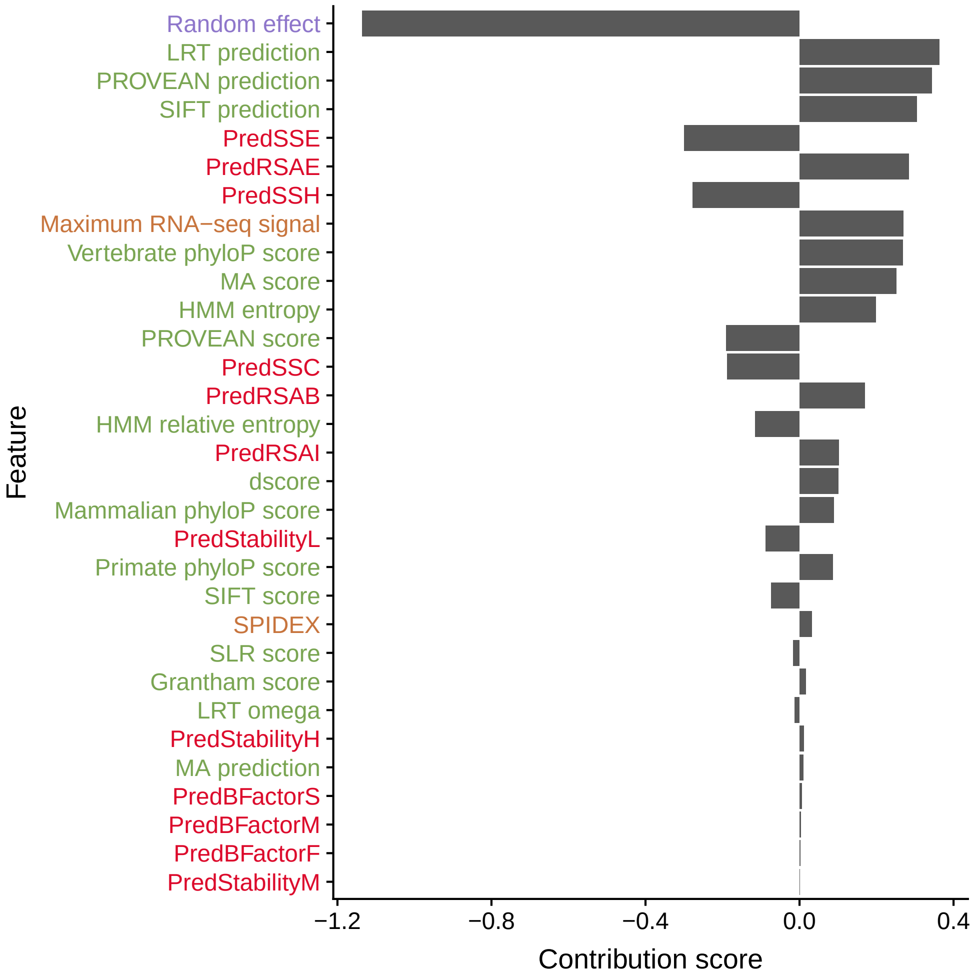


Supplemental Figure S5: Feature contribution scores from the linear UNEECON model. A positive contribution score suggests that the corresponding feature is positively correlated with the strength of selection, while a negative contribution score suggests that the corresponding feature is negatively correlated with the strength of selection. The colors of feature names correspond to four groups: gene-level random effect (purple), sequence conservation (green), structural information (red), and regulatory information (orange).
